# A molecularly defined insular cortex --> central amygdala circuit mediates conditioned overconsumption of food

**DOI:** 10.1101/684498

**Authors:** S.A. Stern, L.E. Pomeranz, E.P. Azevedo, K.R. Doerig, J.M. Friedman

## Abstract

Feeding is a complex motivated behavior that is controlled not just by metabolic and homeostatic factors, but also by environmental factors such as emotion and the hedonic nature of the food itself. Yet, little is known about how brain regions involved in cognition and emotion might contribute to overeating, and therefore, obesity. We used a recently developed behavioral task in which learned contextual cues induce feeding even in sated mice to investigate the underlying neural mechanisms. Using viral tracing, molecular profiling and chemo/optogenetic techniques, we discovered that an insular cortex projection to the central amygdala is required for conditioned overconsumption but not homeostatic feeding. The projection neurons express nitric oxide synthase-1 and activation of this population suppresses satiety signals in the central amygdala. The data thus indicate that the insular cortex provides top down control of homeostatic circuits to promote overconsumption in response to learned cues.

**One Sentence Summary:** Nitric oxide synthase-1 neurons in the insular cortex promote overconsumption by projecting to the central amygdala to suppress a homeostatic satiety signal.

---

Increased food consumption leading to obesity now affects more than 30% of Americans, a public health crisis costing in excess of $150 billion per year (*1-2*). Feeding is a complex motivated behavior that is coordinated by homeostatic (e.g. alteration of energy stores) and non-homeostatic factors (e.g. hedonic, emotional and cognitive cues) (*3-7*). While the neural circuits mediating homeostatic feeding have been extensively delineated, less is known about the neural mechanisms by which learned environmental cues can regulate food consumption. Environmental cues are known potentiators of feeding in both humans and rodents *(8-11)*. To study cue-induced overeating, we have recently described a behavioral paradigm in mice that uses appetitive Pavlovian conditioning to induce overconsumption in response to contextual cues (*12*). In this experiment, fasted mice are placed in a novel environment where food is available and they are allowed to eat until they are satiated. We found that returning these mice to this same environment at a later time led to increased food consumption relative to controls even when the mice were fed. The data indicated that the mice had associated the cues in this environment with a prior experience of hunger (and food availability) leading the mice to continue to eat even when they were satiated. Thus mice previously given food in a novel context when hungry (Ctx+) ate more food when returned to this environment than control mice (Ctx-) even when they were satiated (Fig. 1A, left panel and right panel, blue bars). We developed this paradigm of conditioned overconsumption in order to define those neural circuits that can induce food consumption even in the absence of hunger.

**Fig. 1.**
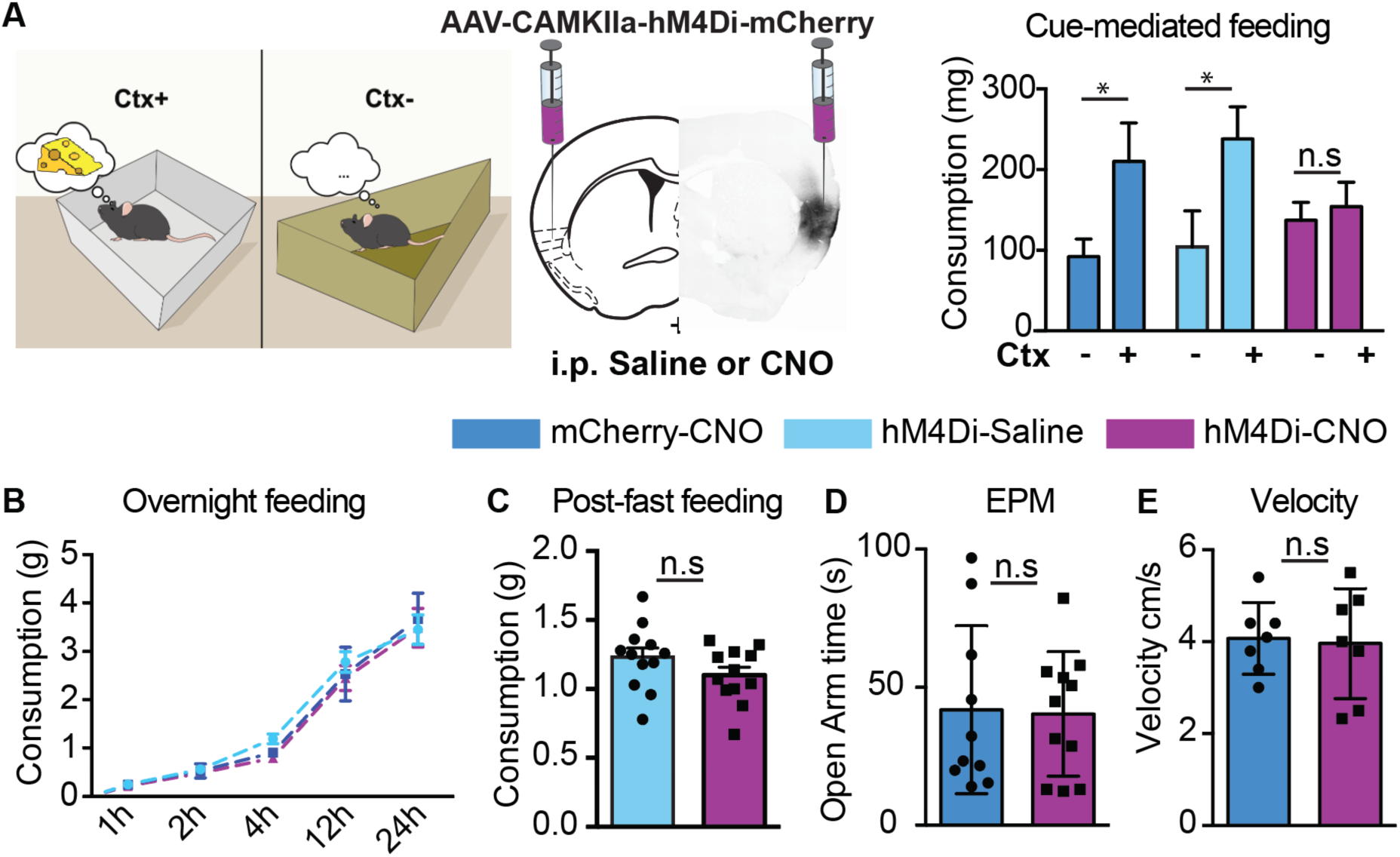
The insular cortex is required for conditioned overconsumption, but not homeostatic feeding. (A) Schematic of learned overconsumption test (left). Mice injected with AAVs encoding hM4Di or control vector (middle) in the insular cortex were tested for their overconsumption response in either the Ctx+ or Ctx- (right). Consumption (measured in milligrams, mg) of mice injected with either CNO (mCherry: dark blue bars, n=4; hM4Di: purple bars, n=10) or saline (hM4Di: light blue bars, n=5) (two-way RM ANOVA, F(1,16)=14.04, **p=0.0018 followed by Holm’s-Sidak multiple comparisons). (B) Food consumption, measured in grams (g) over a 24 hour period (Overnight feeding, n=4-8). (C) Food consumption, measured in g, for 1 hour following an overnight fast (Post-fast feeding, n=12). (D) Time, measured in seconds, s, spent in the open arms of an elevated plus maze (EPM) (n=10-11). (E) Velocity, measured in centimeters transversed per s (cm/s) (n=7).

We recently found that the insular cortex, a region previously implicated as a potential site of integration for motor outputs regulating feeding (*13*), is activated by contextual cues associated with overconsumption and that pharmacological inhibition of this region prevents conditioned overconsumption (*12*). In this report, we began by testing whether inhibition of projection neurons in the insular cortex would prevent learned overconsumption using a CAMKIIa promoter to express an inhibitory DREADD (Designer Receptors Exclusively Activated by Designer Drugs) in the insular cortex (Fig. 1A, middle panel). To that end, we targeted an AAV expressing CAMKIIa-hM4Di to the posterior part of the anterior insular cortex on a rostral-caudal axis (Fig S1). We found that inhibition of the insular cortex projection neurons prior to measuring food intake in trained, fed mice placed in Ctx+ blocked the increase in food intake of trained mice expressing the inhibitory DREADD but treated with saline in the test for overconsumption (Fig 1A, right panel, light blue bars *p<0.05; purple bars, n.s..) There was also a significant decrease in feeding in the conditioned overconsumption test compared to control mice injected with a cre-dependent mCherry virus and given CNO. (Figure 1A, right panel, dark blue bars *p<0.05; purple bars, n.s., *p<0.05). In contrast, inhibition of the insular cortex had no effect on homeostatic feeding, as there was no difference in the 24 hour food intake of naïve mice kept in their home cage (Fig 1B; Overnight feeding, n.s.), nor did it affect food intake after refeeding (Fig 1C; Post-fast Feeding, n.s.). Inhibition of the projection neurons in the insular cortex also had no effect on anxiety behavior in an elevated plus maze task or general locomotor activity (Figure 1D-E, n.s.). These results indicate that glutamatergic projection neurons in the insular cortex mediate conditioned overconsumption, but do not regulate homeostatic feeding (*14,15*).

To identify the neural sites responsible for this effect of insular cortex inhibition, we mapped the projection sites from the insular cortex by injecting an AAV expressing CaMKIIa-mCherry into the insular cortex and visualizing the sites of mCherry labeled terminals. We identified several projection sites from the insular cortex (Figure S2) including a dense projection to the central amygdala (CeA) (Fig. 2A), similar to that previously reported for the anterior insular cortex (*16-19*). We further confirmed this projection anatomically by tracing from the CeA using a retrograde-AAV expressing GFP (Fig. 2B). We tested whether this projection was functional by expressing an activating hM3Dq in IC projection neurons and found increased Fos expression in the CeA after CNO treatment relative to controls (Fig. 2C, p<0.001). We further tested the functional role of inhibiting this projection on the conditioned overconsumption response by injecting the CAMKIIa-hM4Di AAV into the insular cortex and injecting CNO locally at the CeA terminals. We found that inhibition of the insular cortex → CeA projection specifically blocked learned overconsumption (Fig. 2D, blue bars *p<0.05; purple bars, n.s.), without having any effect on refeeding after an overnight fast (Fig. 2E), anxiety or locomotion (Fig S3). We also performed a real-time place preference (RTPP) test and found that inhibition of the insular cortex → CeA projection using cre-mediated retrograde expression of Arch3.0, led mice to avoid the light-paired side of the RTPP chamber (Fig. 2F), indicating that inhibition of this projection is aversive. This suggests that inhibition of this projection prevents overconsumption by decreasing the rewarding aspects of the cue-food association.

**Figure 2:**
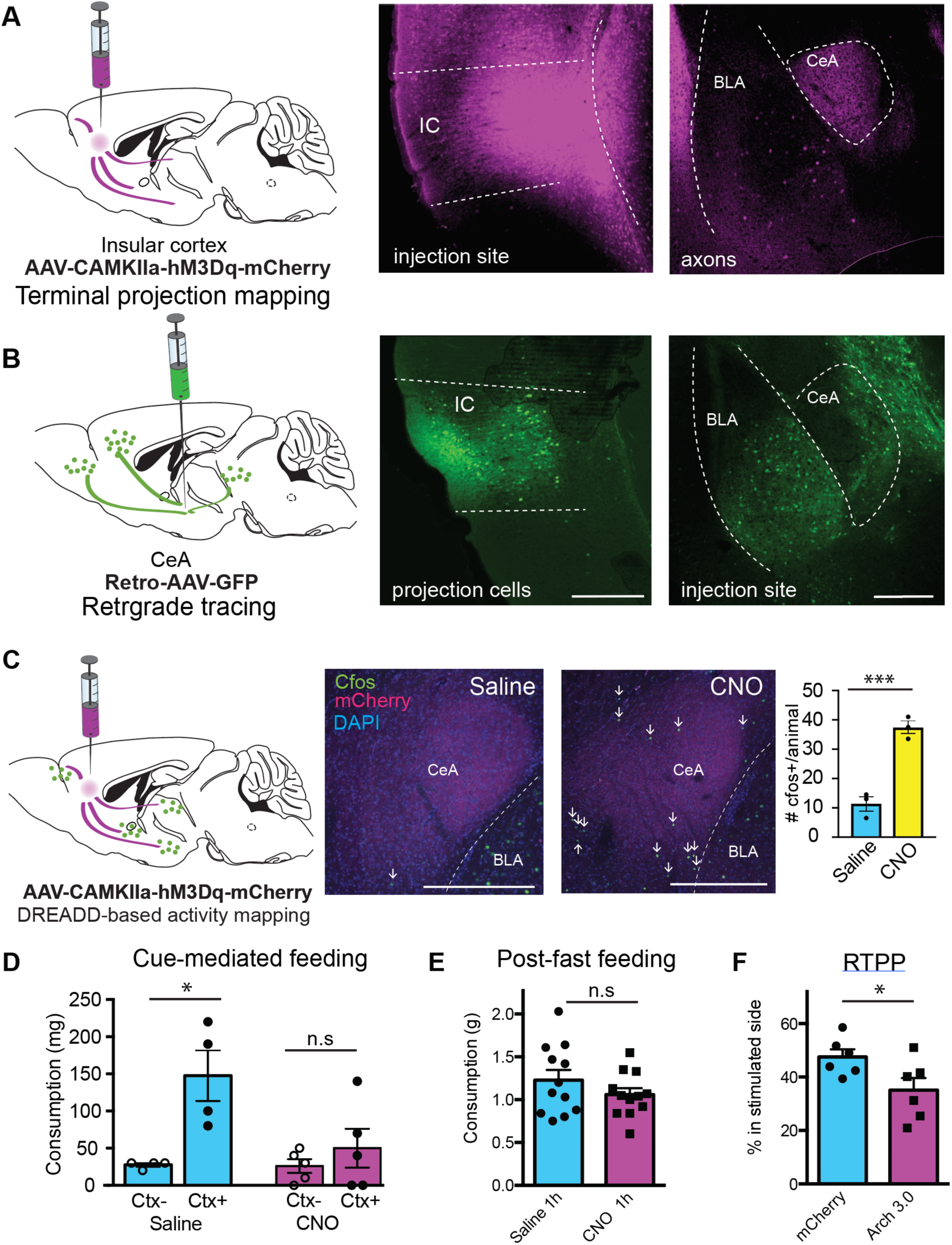
(A) Representative images of terminal projection mapping (left) from the insular cortex injection site (middle) and axon terminals in the central amygdala (right). (B) Representative image of retrograde tracing (left) from the central amygdala injection site (right) and cell bodies in the insular cortex (middle). (C) Schematic of DREADD based activity mapping (left) Representative images of the number of cfos+ cells in the central amygdala in mice injected with saline (right) compared to CNO (middle). Quantification of the number of cfos+ cells, averaged for each animal (right, Nested t-test: t=6.98, df =11, ****P<0.0001). (D) Consumption, measured in mg, of mice expressing hM4Di in the insular cortex and injected with CNO or saline into the CeA prior to the overconsumption test (n=4-5, two-way ANOVA, CNO: F(1,14)=5.2, *P=0.04, CS: F(1,14)=11.0, **P=0.005, Interaction: F(1,14)=4.9, *P=0.04), followed by Sidak’s multiple comparison test. (E) Consumption, measured in g, of mice expressing hM4Di in the insular cortex and injected with CNO or saline into the CeA prior to refeeding (n=11-12). (F) The percentage of time spent (%) in the stimulated side of an RTPP chamber in mice expressing Arch3.0 in insular cortex → CeA projection neurons (n=6, unpaired t-test, t=2.3, df=10, *P=0.04). Scale bars, 400µm.

We next set out to identify the neuronal cell type(s) in the insular cortex responsible for this effect using retro-TRAP, a method that enables molecular profiling of projection neurons (*20*). We injected the retrograde viral tracer, CAV-GFP, into the CeA of SYN-NBL10 mice, which express a camelid nanobody to GFP fused to the L10 subunit of the ribosome in neurons. We then dissected the insular cortex, and performed an immunoprecipitation (IP) to pull down mRNAs associated with the GFP-bound fraction, which correspond to the CeA projection neurons (Fig. 3A). The enrichment of each gene was calculated as the number of reads in the precipitated RNA relative to the total (IP/input [INP]) and we further assessed the validity of this assay in our system with qPCR. *Gfp*, a positive control, was highly enriched in the IP fraction, whereas glial markers *Gfap* and *Mal* were depleted (Fig. 3A, bottom left). We then generated a dataset of significantly enriched and depleted genes (q-value <0.05), and identified molecular markers that were significantly enriched in the insular cortex → CeA projection relative to the entire population of insular cortex neurons (Fig. 3B). We found ∼2500 significantly enriched genes out of ∼18,000 that were detectable, and ∼2000 of those had an enrichment of 2-fold or greater (Table S1).

**Fig 3:**
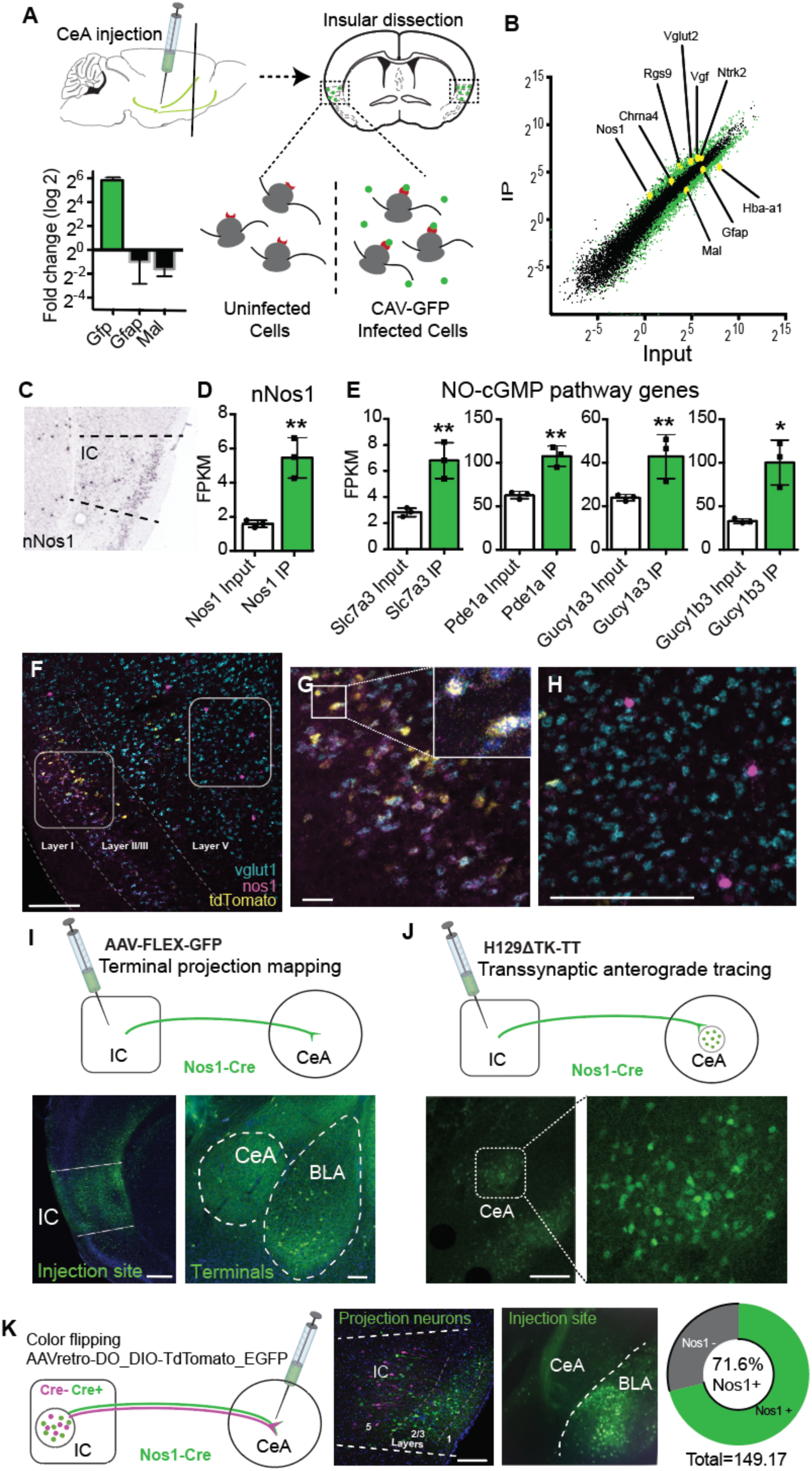
Nos1 is a marker for insular cortex to central amygdala projection neurons. (A) Experimental scheme for retro-TRAP experiment. Syn-NBL10 mice are injected with CAV-GFP into the central amygdala and polysomes bound by GFP in the insular cortex are precipitated out, representing the insular cortex → CeA projection neurons. Lower left, the fold change of positive (*Gfp*, green) and negative (*Gfap* and *Mal*, black) controls run by qPCR. (B) Plot depicting the average IP value and average Input values (log2) of all genes detected during sequencing. Genes significantly enriched and depleted with a q-value <0.05 are represented in green. Selected genes on interest are highlighted in yellow. (C) Representative *in situ hybridization* image depicting Nos1 expression in the insular cortex (courtesy of Allen Brain Atlas). (D) Enrichment of Nos1 FPKM in the IP fraction compared to the Input (n=3, unpaired t-test, t=5.59, df=4, **P=0.005). (E) Enrichment of Nitric Oxide – cGMP pathway genes in the IP fraction compared to the Input (n=3, unpaired t-tests: *Slc7a3* t=4.76, df=4, **P=0.009; *Pde1a* t=6.28, df=4, **P=0.003; *Gucy1a3* t=3.2, df=4, *P=0.03; *Gucy1b3* t=4.49, df=4, *P=0.011). (F) Representative *In situ hybridization* in the insular cortex of vGlut1 (cyan), Nos1 (magenta) and tdTomato (yellow) from Nos1-cre mice injected with AAVretrograde-mCherry into the CeA. Scale bar, 200µm. Left inset of F is magnified in G, depicting layer 2-3 neurons expressing all three markers. Scale bar, 40µm The right inset is magnified in H depicting Nos1 neurons that do not express vGlut1. Scale bars, 200µm. (I) Representative images of terminal projection mapping (left) from the insular cortex injection site of Nos1-Cre mice (middle) and axon terminals in the CeA (right). Scale bar, 400µm. (J) Representative images of central amygdala neurons (right) of mice injected with the transsynaptic anterograde tracer H129ΔTK-TT from insular cortex of Nos1-cre mice. Scale bar, 400µm. (K) Representative images of Nos1-Cre mice injected with the color-flipping Cre-dependent retrograde AAV from the CeA (left). Nos1+ cells (green) and Nos1-cells (magenta) are labeled in the insular cortex (middle). Scale bar, 150µm. Quantification of the percentage of Nos1+ neurons in the CeA projecting population (right).

One of the genes that was significantly enriched (∼3-fold) in our dataset was nNos1 (nitric oxide synthase-1, Nos1; Fig. 3C-D, **P<0.01). We also found significant enrichment of Slc7a3, Pde1a, Gucy1a3 and Gucy1b3 (*23*), all of which are known to be co-expressed in Nos1 neurons (Fig. 3E, *P<0.05, **P<0.01). While Nos1 neurons within the hypothalamus have been implicated in feeding behavior (*21,22*), less is known about the function of Nos1 neurons in cortex. However, in cortex, Nos1 is primarily known as a marker for GABAergic interneurons populations (*23*), though other data have shown that there is also a separate population of Nos1 projection neurons (*24*), and that some excitatory pyramidal cells can express Nos1 (*25*). In our dataset, there was a depletion of molecular markers for inhibitory interneurons including Sst, Pvalb, Vip, Npy, Cort, Calb2, and Cck (Figure S4A), but enrichment of markers such as Rgsp, Syt4, Hhip, Trpc5 and Tacr1, which have also been previously reported to be expressed in Nos1 projection neurons (*23*) (Figure S4B). These findings suggested that Nos1 marks a specific population of neurons that project from the insular cortex → CeA, but it was unclear if they were GABAergic or glutamatergic. To confirm their neurotransmitter identity, we performed *in situ hybridization* of brains from Nos1-cre mice after injection with a retrograde cre-dependent AAV expressing tdTomato into the CeA to label Nos-1 neurons in the insular cortex that project to the CeA (Figure S5A). We found that a population of insular cortex Nos1 neurons in layers 2 and 3 project to the CeA (and express tdTomato) and exclusively express vGlut1 (Figure 3F-H). In contrast, a separate population of neurons expressing high levels of Nos1 in layer 5, and which did not express tdTomato (and thus did not project to CeA) did not express vGlut1 and likely correspond to GABAergic interneurons (*25*) (Fig. 3F-H). Indeed, we find that Nos1 neurons expressing vGAT do not co-express tdTomato and therefore do not project to the CeA (Figure S5B-D). These data indicate that a specific population of glutamatergic Nos1 neurons in the insular cortex project to the central nucleus of the amygdala.

We further confirmed that a subset of insular Nos1 neurons project to the CeA using other methods for neuronal tracing: we first examined the projection in the anterograde direction by injecting an AAV-FLEX-mCherry (Fig. 3I) and a transsynaptic anterograde tracing herpes simplex virus H129ΔTK-TT virus (*26*) into the insular cortex of Nos-1 cre mice (Fig 3J). These data confirmed that Nos1 neurons project to CeA. To quantitate these data, we injected a color flipping retrograde AAV which expresses GFP in the presence of Cre, and tdTomato in its absence, into the CeA of Nos1-cre mice and analyzed expression of GFP vs. tdTomato in the insular cortex. We found that Nos1 neurons comprise ∼71.6% of the total number of IC neurons that project to the CeA (Fig. 3K). Consistent with the previous data (see Fig 3F-H), nearly all Nos1 → CeA projection neurons (∼90%) were located in more superficial layers 2-3 of the insular cortex, and were sparse in deeper layers (Fig. 3F, 3K).

We then tested whether the insular cortex Nos1 neurons are functionally required for the conditioned overconsumption response. In line with our previous results, inhibition of the insular cortex Nos1 neurons prior to the overconsumption test (i.e; food intake of fed mice in Ctx+ vs. Ctx-) completely blocked the overconsumption exhibited by control Ctx+ mice (Fig. 4A), without affecting homeostatic feeding, locomotion or anxiety (Figure S6). In these studies, we inhibited Nos1 neurons by injecting a cre-dependent AAV expressing hM4Di into the insular cortex of Nos1-cre mice.

**Fig. 4:**
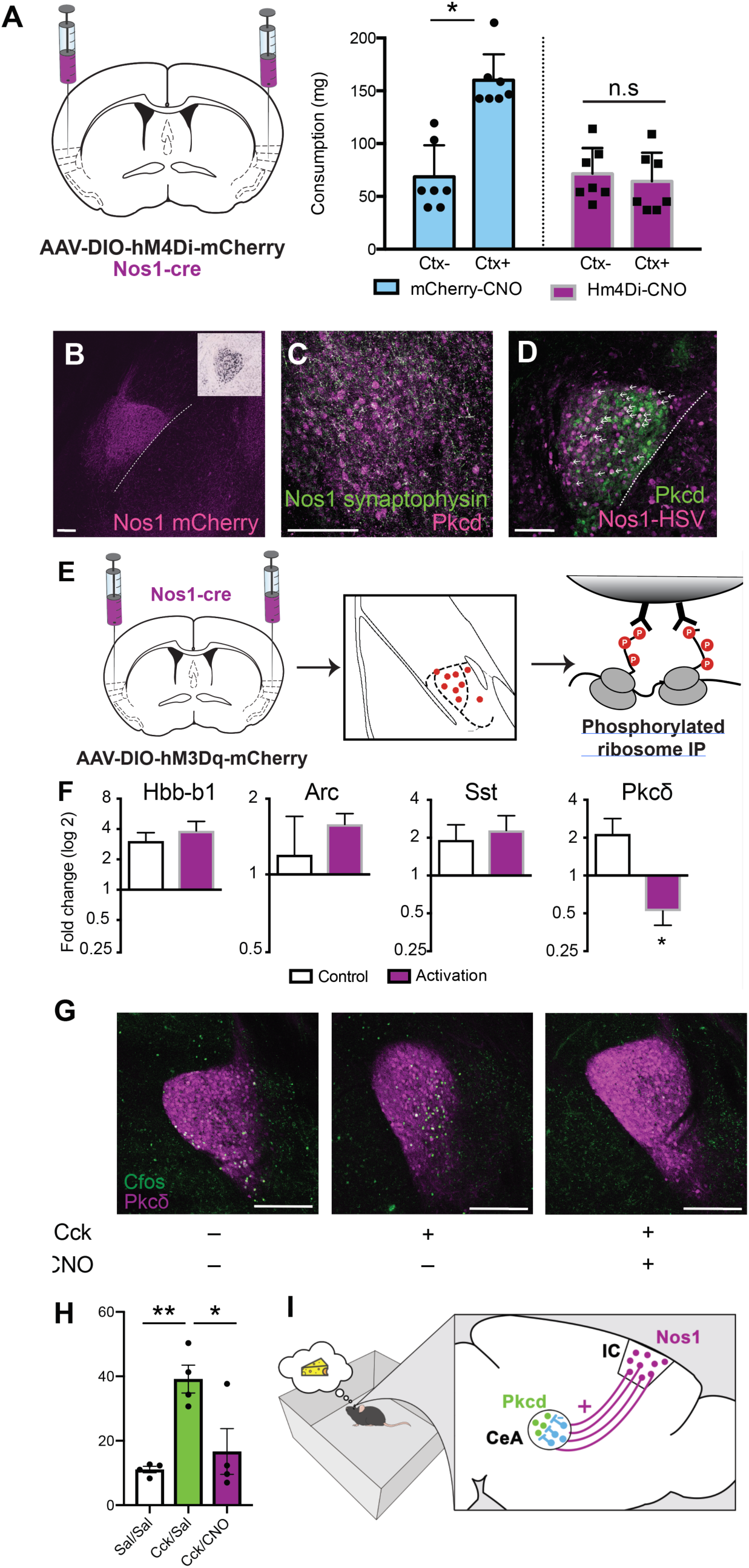
Insular cortex Nos1 neurons are required for conditioned overconsumption and suppress central amygdala satiety signals. (A) Nos1-cre mice injected with AAVs encoding hM4Di or control vector (left) in the insular cortex were tested for their overconsumption response in either the Ctx+ or Ctx- (right). Consumption, measured in mg, of Nos1-cre mice expressing hM4di (purple bars) or mCherry (blue bars) and injected with CNO prior to the overconsumption test. (Interaction: F(1,12)=4.34, P=0.059; CS: F (1, 12) = 3.173, P=0.1002; DREADD: F(1, 12) = 2.546, P=0.1366, with Sidak’s multiple comparisons post-hoc test, *P<0.05). (B) Representative image of Nos1 terminals in the CeA (magenta). Inset depicts Pkcδ in the CeA (Allen Brain Atlas). (C) Representative image of synaptophysin labeled terminals (green) from the insular cortex of Nos1-cre mice, localized in the same location as Pkcδ neurons (magenta) in the CeA. (D) Representative image of anterograde tracing (magenta) revealing insular cortex Nos1 neurons that are synaptically connected to Pkcδ neurons (green) in the CeA (white arrows). (B-D) Scale bars, 150µm. (E) Phospho-TRAP experimental scheme. Insular cortex Nos1 neurons were activated with hM3Dq DREADDs and pS6+ neurons in the CeA were immunoprecipitated out one hour later. (F) Fold-change determined by qPCR of quality control (Arc, Hbb-b1) and CeA, (Sst, Pkcδ) markers. (Paired t-test: t=4.822, df=2, *P=0.04) (G) Representative images depicting co-expression of Pkcδ (magenta) and cfos (green) in Nos1-cre mice injected with AAV-DIO-hM3Dq-mCherry into the insular cortex and injected with saline+saline (left), Cck+Saline (middle) or Cck+CNO (right). Scale bars, 250µm. (H) Quantification of G (nested One-Way ANOVA, **p=0.0039 followed by Tukey’s Multiple Comparison Test, **p<0.01, *p,0.05). (I) Model of learned overconsumption in which mice are placed in the food-associated context, leading to Nos1 inhibition of satiety signals in Pkcδ neurons in the CeA.

The terminals of the insular cortex Nos1 projection neurons are localized in the lateral portion of the CeA which is also where anorexigenic Pkcδ (Protein Kinase C, delta) neurons reside (*16*) (Fig. 4B). These Pkcδ neurons are activated by food intake as part of a satiety response. We therefore hypothesized that insular cortex Nos1 neurons might promote conditioned overconsumption by inhibiting Pkcδ neurons in the CeA. Consistent with this, injection of an AAV expressing a Flex-GFP-synaptophysin construct into the insular cortex of Nos1-cre mice showed synaptophysin expression in nerve terminals in the same, lateral region of the CeA as Pkcδ neurons (Fig. 4C). Furthermore, an injection of the transsynaptic anterograde tracer, H129ΔTK-TT into the insular cortex of Nos1-cre mice labeled Pkcδ neurons in the CeA (Fig. 4D, white arrows). This virus serially infects neurons across synapses and thus labels both direct and indirect targets of the Nos-1 neurons. Because the Nos1 neurons are glutamatergic and activation of the Pkcδ neurons would be expected to decrease food intake rather than leading to overconsumption, we hypothersized that Pkcδ neurons are not a direct target of insular cortex Nos1 neurons. Rather, we hypothesized that the Nos1 neurons indirectly inhibit the Pkcδ neurons, perhaps by activating a local GABAergic neuron than in turn inhibits the Pkcδ neurons.

To test whether insular cortex Nos1 neurons modulate CeA Pkcδ neuron function we used phospho-TRAP (phospho-translating ribosome affinity purification) (*27*) to show depletion of Pkcδ in the CeA after activation of the insular cortex Nos1 neurons. Depletion of markers using phospo-TRAP is an indicator of neural inhibition. This approach was used because Fos is a marker for neural activation and decreased levels of pS6 is a biochemical marker of neural inhibition. To that end, we activated insular cortex Nos1 neurons infected with an AAV expressing cre-dependent hM3Dq and immunoprecipitated polysomes using a phospho-S6 (pS6) antibody and analyzed the enrichment and depletion of specific mRNAs in the CeA (Fig. 4E). As expected, qPCR revealed enrichment for *Arc*, a marker of activation, and *Hbb-b1* which is a control marker highly enriched with pS6-expressing cells (*28*) (Fig. 4F). We then analyzed the data using a set of lateral CeA markers (*29,30*) and found that only the gene for Pkcδ, *prkcd*, was depleted after Nos1 neural activation (∼4 fold compared to saline-injected controls, *p<0.05). No other CeA markers that we tested were depleted indicating that the *prkcd* population is inhibited following Nos1 activation (Fig. 4F, Fig S7). To further confirm that Pkcδ neurons are indirectly inhibited by insular cortex Nos1 projection neurons, we injected mice with the satiety-inducing cholecystokinin (Cck) i.p., which leads to activation of Pkcδ neurons and thus, induces the expression of Cfos in these neurons (*16,29*), while simultaneously activating Nos1 neurons using hM3Dq activatory DREADDs. We found that activating insular cortex Nos1 neurons completely blocked the Cck-induced increase in Cfos in Pkcδ neurons (Fig. 4G-H). Because most CeA neurons are GABAergic, this suggests that insular cortex Nos1 neurons project to a subset of CeA neurons which in turn inhibit Pkcδ neurons to promote overconsumption (Figure 4I).

In summary, we find that glutamatergic insular cortex Nos1 neurons provide top-down control of learned overconsumption via a projection to central amygdala. This provides a molecularly defined neural circuit mechanism through which environmental cues can promote food intake and which may, over time, lead to increased body weight. These findings are consistent with prior reports showing that the insular cortex is required for Pavlovian cue-food associations (14,31). The insular cortex is also required for conditioned-taste aversion (*32*), and these data together with ours provides evidence that this brain region can regulate food intake in response to specific learned cues in either direction. It will thus be interesting to establish whether the effect of conditioned taste aversion is mediated by Nos1 neurons or a different neural population. The central amygdala is known to regulate homeostatic feeding and our data further suggests that learned responses regulating feeding are controlled by top down inputs from cortex to subcortical pathways that also play a more general role in the control of homeostatic feeding. These studies thus provide a potential framework for understanding how prior experience can regulate a basic behavior.

It will now be important to establish how sensory information is decoded by the insular cortex and linked to neural pathways associated with the prior experience of hunger and food availability. The insular cortex is known to receive a large number of sensory inputs that are relevant for feeding including taste as well as visceral signals. The emotional valence of the prior experience is also relevant and this could also involve reciprocal connections with the amygdala. For example, a previous study showed that a circuit from the basolateral amygdala to insular cortex is required for learning about cue-food associations under periods of food deprivation (14). This paradigm is comparable to the cue-food associations established during the training sessions in the conditioned overconsumption task we developed.

Overall, these studies provide new insight into how the brain links diverse environmental stimuli to modulate innate behavioral responses under specific conditions. Our studies invoke the insular cortex as playing an important role in these conditioned responses. Consistent with this, human imaging studies have suggested that the insular cortex activity is altered in patients with obesity *(33-35)*. These findings thus provide potential insights into how specific environmental cues in human (such as advertising or eating in front of the television) can lead to increased food intake in specific circumstances. This work is thus highly relevant to understanding how environmental factors can regulate food intake potentially leading to obesity.

## Materials and Methods

### Animals and breeding

C57BL/6J and Nos1-cre mice were obtained from Jackson Laboratories. SYN-NBL10 mice were generated in our laboratory (Ekstrand et al. 2014). Mice were housed according to the guidelines of the Rockefeller’s University Animal Facility. Males and females at age 8-20 weeks were used throughout this study and all animal experiments were approved by the Rockefeller University IACUC, according to NIH guidelines. Male mice were used for behavioral experiments, but both male and female mice were used for anatomical studies, including TRAP. Mice were kept on a 12-h/12-h light/dark cycle (lights on at 7:00 a.m.) and had access to food and water ad libitum, except when noted otherwise.

### Viral vectors

All AAVs used in this study were purchased from UNC Vector Core or Addgene, except where noted. AAV5-CaMKIIa-hM4Di-mCherry, AAV5-DIO-hM4Di-mCherry, AAV-DIO-eNphr3.0-EGFP were used for inhibition studies. AAV5-CaMKIIa-hM3Dq-mCherry or AAV5-DIO-hM3Dq-mCherry were used for activation studies. Control AAV-CaMK2a-mCherry or AAV-CAMK2a-were used for comparison in control WT mice. AAV-DIO-mCherry or AAV-DIO-EGFP viruses were used for comparison in Cre+ littermate control mice. pAAV-Ef1a-DO_DIO-TdTomato_EGFP-WPRE-pA (color-switching virus), AAVretro-hSyn-GFP and AAVretro-FLEX-tdTomato were used for tracing studies. The retrograde tracer CAV-GFP was obtained from Montpelier and used for retro-TRAP experiments. H129ΔTK-TT was a gift from David Anderson.

### Pharmacology

CNO (Sigma) were dissolved in saline with 0.1% DMSO and injected either intraperitoneally (i.p) at 5 mg/kg or through a cannula at 500uM. Behavioral tests were performed 30-40 minutes after drug injection. CCK (Tocris) was injected i.p. at 5 µg/kg. Mice were euthanized 1 hour following CCK injection for cfos analysis.

### Stereotaxic Injections

Mice were anesthetized with isoflurane, placed in a stereotaxic frame (Kopft Instruments) and bilaterally injected in either the insular cortex or the central amygdala using the following coordinates relative to bregma: Insular Cortex (AP: +0.26; ML: 3.85; DV: −3.9); Central Amygdala(AP: −1.4; M/L: 2.8; DV: −4.72) (Paxinos). A total of 200-400 nL of virus at high titer concentrations (at least 10^11^) were injected per site at a rate of 100 nL/min. For pharmacological experiments, cannula were implanted in the central amygdala at the same coordinates as above. For optogenetics experiments, Optic fiber implants (Thor Labs) were implanted over the CeA at the following coordinates relative to bregma: AP: −1.4; ML: 2.8; DV: −4.4 (Paxinos). Implants were inserted slowly and secured to the mouse skull using two layers of Metabond (Parkell Inc) followed by a layer of dental cement. Mice were single housed and monitored in the first weeks following optic implantation. For all surgeries, mice were monitored for 72h to ensure full recovery and two weeks later mice were used in experiments and were sacrificed following behavioral studies to confirm viral expression and fiber placement using immunohistochemistry.

### Context-Induced Feeding

Prior to habituation mice were given 5-10 chocolate flavored 20mg Precision Pellets during each day of handling to prevent neophobia. Mice were habituated to two different contexts. Contexts were easily distinguishable based on shape, size, floor texture and were in different rooms. Habituation consisted of 20 min exposure to a novel context. Mice were returned to their home cages after habituation, and one hour later, all cages were changed to prevent food dust on the floor from being consumed and food removed for the next 18-24 hours. The next day mice designated at CS+ were trained in the habituation context, and mice designated at CS-remained in their homecage. Training consisted of 30 min exposure to the context with 2g of Precision Pellets freely available in a food well. Pellets were counted after the training session to determine food consumption. Mice were refed *ad lib* for 24h and then fasted again for 18-24h before a second training session. One hour after the second training session, all mice were returned to ad libitum feeding for 48h, at which time testing was conducted. Testing consisted of both CS+ and CS-mice undergoing a 20 min exposure to the habituation context with 2g of Precision Pellets freely available in a food well. Pellets were again counted after testing to determine food consumption. In these experiments, mice were injected with CNO or saline 1 hour prior to the overconsumption test.

### Overnight Feeding

Mice were injected with CNO or saline 1 hour prior to dark phase onset with pre-measured food available in the food hopper of their homecage. Chow was measured at 1h, 2h 4h and 24h post injection.

### Post-Fast Feeding

Mice were fasted for 24h starting 3-4h after the light-phase onset in their homecage. Mice were subsequently injected with CNO or saline and then refed. Food was measure 1h and 2h after refeeding.

### Elevated Plus Maze

Mice were placed at the center of a cross-shaped, elevated maze in which two arms are closed with dark walls and two arms are open and allowed to explore for 10 min. Mice were injected with CNO (1mg/kg) 1h before testing. After, mice were returned to their homecage and the maze floor was cleaned in between subjects. All subjects were recorded using a camera and behavior (time spent in open and closed arms, distance and velocity) were analyzed using Ethovison 9.0 (Noldus).

### Real Time Place Preference (RTPP)

Mice were habituated to optic patch cables for 3-5 days for ∼3 minutes each. On a baseline day they were first acclimated to the RTPP chamber by placed on the border between two adjoining homogeneous black compartments with white floors. The amount of time spent in each compartment was recorded using video tracking software (Ethovision 9, Noldus) for 10 minutes to ensure that there was no explicit side preference. On the subsequent day, one side was designated as light-paired in which active entry triggered photostimulation (<u>inhibition</u>: constant light; <u>activation</u>: 1 Hz, 10 ms pulse width, 5-10 mW), using lasers controlled by a Mini I-O box from Noldus and a waveform generator (Keysight). Sessions lasted for 20 min and the amount of time spent in each compartment was recorded.

### Retro-TRAP

Mice were injected with CAV-GFP into the CeA and fourteen days later mice were sacrificed by cervical dislocation and the insular cortex was rapidly dissected in ice-cold Buffer B (1xHBSS, 4 mM NaHCO_3_, 2.5 mM HEPES [pH 7.4], 35 mM Glucose) with 100 mg/ml cycloheximide (Sigma). The dissected pieces were pooled in 3 groups of 5-6 brains each and transferred to a glass homogenizer (Kimble Kontes 20), and homogenized in 1.5 ml ice-cold Buffer C (10 mM HEPES [pH 7.4], 150 mM KCl, 5 mM MgCl_2_) with 0.5 mM DTT (Sigma), 80 U/ml RNasin Plus (Promega), 40U/ml Superase-In (Life Technologies), 100 mg/ml cycloheximide, protease inhibitor cocktail (Roche) and 100 ng/ml GFP-Trap Protein (ChromoTek). Tissue samples were homogenized three times at 250 rpm and ten times at 750 rpm on a variable-speed homogenizer (Glas-Col) at 4°C. Homogenates were transferred to microcentrifuge tubes and clarified at 2,000xg for 10 min at 4°C. 140 µl each of 10% IGEPAL CA-630 (NP-40; Sigma) and 1,2-diheptanoyl-sn-glycero-3-phospho-choline (DHPC at 100 mg/0.69 ml; Avanti Polar Lipids) was added to the supernatant. The solutions were mixed and centrifuged again at 17,000xg for 15 min at 4°C. The resulting supernatants were transferred to new tubes and 50 µl of each cleared lysate was mixed with 50 µl Lysis Buffer (0.7 µl β-mercaptoethanol/100 µl Lysis Buffer; Agilent Absolutely RNA Nanoprep Kit) and stored at –80° for later preparation as input RNA. The remaining lysates (approximately 1.5 ml) were used for immunoprecipitation. The beads incubating with GFP antibodies were washed twice in Buffer A with 0.5 mM DTT, 80 U/ml RNasin Plus and 100 mg/ml cycloheximide before the cleared brain lysates were added. The immunoprecipitation was allowed to run at 4°C for 40 min. Beads were washed four times with Buffer D (10 mM HEPES [pH 7.4], 350 mM KCl, 5 mM MgCl_2_, 1% NP40) with 0.5 mM DTT, 80 U/ml RNasin Plus and 100 mg/ml cycloheximide. Before removing the last wash solution the beads were moved to a new tube. After the final wash, RNA was eluted by adding 100 µl Lysis Buffer and purified using the Absolutely RNA Nanoprep Kit (Agilent).

### RNA Sequencing

cDNA was amplified using SMARTer Ultralow Input RNA for Illumina Sequencing Kit and sequenced on an Illumina HiSeq2500 platform. RNA sequencing raw data was uploaded and analyzed using BaseSpace apps (TopHat and Cufflinks; Illumina) using an alignment to annotated mRNAs in the mouse genome (UCSC, Mus musculus assembly mm10). The average immunoprecipitated (IP) and Input value of each enriched and depleted genes with a q-value lower than 0.05 were plotted using GraphPad Prism (GraphPad).

### Phospho-TRAP

The PhosphoTrap profiling experiments were performed according to Knight et al., 2012. Briefly, mice injected with AAV5-DIO-Hm3dq in the insular cortex (coordinates above) were separated into groups of 4-6 mice per group (termed control or activation). Control mice were given a saline injection and Activation mice were given an injection of CNO (3mg/kg). Mice were euthanized 1 hour post-injection, brains were removed, and the central amygdala were dissected on ice and pooled into 3 replicates per group of 4-6 mice each. Tissue was homogenized and clarified by centrifugation. Ribosomes were immunoprecipitated by using 4 ug of polyclonal antibodies against pS6 (Invitrogen) previously conjugated to Protein A-coated magnetic beads (Thermofisher). A small amount of tissue RNA was saved before the immunoprecipitation (Input) and both input and immunoprecipitated RNA (IP) were then purified using RNAeasy Mini kit (QIAGEN) and RNA quality was checked using a RNA PicoChip on a bioanalyzer. RIN values > 7 were used. For qPCR analysis cDNA was prepared with the QuantiTect Reverse Transcription Kit (QIAGEN).

### qPCR analysis

qPCR using predesigned Taqman probes (idtDNA) were used and cDNA was prepared using the QuantiTect Reverse Transcription Kit (Life Technologies). The abundance of these genes in IP and Input RNA was quantified using Taqman Gene Expression Master Mix (Applied Biosystems). Transcript abundance was normalized to beta-actin. Fold of Change were calculated using standards or with the ΔΔCt method if there was not enough material to make standards.

### Immunohistochemistry, quantifications and imaging

Mice were perfused and brains were postfixed for 24h in 10% formalin. Brain slices were taken using a vibratome (Leica), blocked for 1h with 0.3% Triton X-100, 3% bovine serum albumin (BSA), and 2% normal goat serum (NGS) and incubated in primary antibodies for 24h at 4°C. Then, free-floating slices were washed three times for 10 min in 0.1% Triton X-100 in PBS (PBS-T), incubated for 1h at room temperature with secondary antibodies, washed in PBS-T and mounted in Vectamount with DAPI (Southern Biotech). Antibodies used here were: anti-cfos (1:500; Cell Signaling), anti-mCherry (1:1000; Abcam), anti-GFP (1:1000, Abcam), anti-Pkcd (1:500, Millipore) goat-anti-rabbit (Alexa 488 or Alexa 594, Alexa646 1:1000; Thermo Scientific), goat anti-chicken Alexa488, Alexa594, Alexa 647 (1:1000; Thermo Scientific). Images were taken using an LSM780 confocal (Zeiss) and images were processed using ImageJ software (NIH). Cfos counts were conducted for 2-3 sections / animal with a n=2-4 animals / group.

### Fluorescent In Situ Hybridization

For examination of gene expression, tissue samples of mice injected with AAVretro-FLEX-tDTomato underwent single molecule fluorescent in situ hybridization (smFISH). Isoflurane anesthetized mice were decapitated, and brains were harvested and flash frozen on aluminum foil on dry ice. Brains were stored at –80°C. Prior to sectioning, brains were equilibrated to –16°C in a cryostat for 30 min. Brains were cryostat sectioned coronally at 20 µm and thaw-mounted onto Superfrost Plus slides (25×75 mm, Fisherbrand). Slides were air-dried for 60 to 90 min prior to storage at –80°C. smFISH for all genes examined— *Nos1* (Cat# 437691), *mCherry* (Cat# 431202), Vglut1 (Cat# 416631) and Vgat (Cat# 319191) - was performed using RNAscope Fluorescent Multipex Kit (Advanced Cell Diagnostics) according to the manufacturer’. Slides were counterstained for the nuclear marker DAPI using Vectashield mounting medium with DAPI (ThermosFisher). Sections were imaged using an LSM780 confocal (Zeiss) and processed using ImageJ software.

### Statistics

All results are presented as mean ± s.e.m. and were analyzed with Prism software. No statistical methods were used to predetermine sample sizes, but our sample sizes are similar to those reported in previous publications. Normality tests and F tests for equality of variance were performed before choosing the statistical test. Unless otherwise indicated, statistics were based on two-tailed unpaired t tests or Mann–Whitney U tests (for datasets that were not normally distributed) for two-group comparisons. P < 0.05 was considered significant (*P < 0.05, **P< 0.01, ***P < 0.001). Animals in the same litter were randomly assigned to different treatment groups and blinded to experimenters in the various experiments. Injection sites and viral expression were confirmed for all animals. Mice showing incorrect injection sites or optic fiber placement were excluded from the data analysis.

### Data availability

The data that support the findings of this study are available from the corresponding author upon reasonable request.

## Acknowledgments

We thank Ravi Tolwani and the staff of the Comparative Biosciences Center, Connie Zhao and the staff at the Genomics Resource Center and Alison North and the staff at the Bioimaging Resource Center at Rockefeller University for technical assistance. We thank David Anderson for providing the HSV-lsl-tdTomato virus. This work was funded by an F32DK107077 and a NARSAD Young Investigator Award (S.A.S) and the JPB Foundation (J.M.F).

## Author contributions

S.A.S and J.M.F conceived the study. S.A.S designed and conducted the experiments and analyzed data. K.R.D. assisted with immunohistochemistry and behavioral experiments, L.E.P. conducted anterograde tracing, E.P.A assisted with TRAP studies, organizing figures and critical discussions. S.A.S and J.M.F secured funding and wrote the manuscript with the input of all authors.

## Data and materials availability

All data is available in the main text or the supplementary materials, and is available from the corresponding author upon reasonable request.

## References and Notes

1. R. T. Hurt, C. Kulisek, L. A. Buchanan, S. A. McClave, The obesity epidemic: challenges, health initiatives, and implications for gastroenterologists. Gastroenterol. Hepatol. 6, 780–792 (2010).

2. R. A. Hammond, R. Levine, The economic impact of obesity in the United States. Diabetes Metab. Syndr. Obes. 3, 285–295 (2010).

3. C. B. Saper, T. C. Chou, J. K. Elmquist, The need to feed: homeostatic and hedonic control of eating. Neuron. 36, 199–211 (2002).

4. H.-R. Berthoud, Interactions between the “cognitive” and “metabolic” brain in the control of food intake. Physiol. Behav. 91, 486–498 (2007).

5. M. Lutter, E. J. Nestler, Homeostatic and hedonic signals interact in the regulation of food intake. J. Nutr. 139, 629–632 (2009).

6. H.-R. Berthoud, H. Münzberg, C. D. Morrison, Blaming the Brain for Obesity: Integration of Hedonic and Homeostatic Mechanisms. Gastroenterology. 152, 1728–1738 (2017).

7. M. A. Rossi, G. D. Stuber, Overlapping Brain Circuits for Homeostatic and Hedonic Feeding. Cell Metab. 27, 42–56 (2018).

8. H. P. Weingarten, Conditioned cues elicit feeding in sated rats: a role for learning in meal initiation. Science. 220, 431–433 (1983).

9. L. L. Birch, L. McPhee, S. Sullivan, S. Johnson, Conditioned meal initiation in young children. Appetite. 13, 105–113 (1989).

10. C. E. Cornell, J. Rodin, H. Weingarten, Stimulus-induced eating when satiated. Physiol. Behav. 45, 695–704 (1989).

11. G. D. Petrovich, Forebrain circuits and control of feeding by learned cues. Neurobiol. Learn. Mem. 95, 152–158 (2011).

12. S. A. Stern, K. R. Doerig, E. P. Azevedo, E. Stoffel, J. M. Friedman, Control of non-homeostatic feeding in sated mice using associative learning of contextual food cues. Mol. Psychiatry (2018), doi:10.1038/s41380-018-0072-y.

13. C. A. Pérez et al., Molecular annotation of integrative feeding neural circuits. Cell Metab. 13, 222–232 (2011).

14. Y. Livneh et al., Homeostatic circuits selectively gate food cue responses in insular cortex. Nature. 546, 611–616 (2017).

15. B. A. Baldo, R. C. Spencer, K. Sadeghian, J. D. Mena, GABA-Mediated Inactivation of Medial Prefrontal and Agranular Insular Cortex in the Rat: Contrasting Effects on Hunger-and Palatability-Driven Feeding. Neuropsychopharmacology. 41, 960–970 (2016).

16. H. Cai, W. Haubensak, T. E. Anthony, D. J. Anderson, Central amygdala PKC-δ(+) neurons mediate the influence of multiple anorexigenic signals. Nat. Neurosci. 17, 1240–1248 (2014).

17. A. M. Douglass et al., Central amygdala circuits modulate food consumption through a positive-valence mechanism. Nat. Neurosci. 20, 1384–1394 (2017).

18. H. C. Schiff et al., An Insula-Central Amygdala Circuit for Guiding Tastant-Reinforced Choice Behavior. J. Neurosci. 38, 1418–1429 (2018).

19. M. Venniro et al., The Anterior Insular Cortexfi→Central Amygdala Glutamatergic Pathway Is Critical to Relapse after Contingency Management. Neuron. 96, 414–427.e8 (2017).

20. M. I. Ekstrand et al., Molecular profiling of neurons based on connectivity. Cell. 157, 1230–1242 (2014).

21. A. K. Sutton et al., Control of food intake and energy expenditure by Nos1 neurons of the paraventricular hypothalamus. J. Neurosci. 34, 15306–15318 (2014).

22. R. L. Leshan, M. Greenwald-Yarnell, C. M. Patterson, I. E. Gonzalez, M. G. Myers Jr, Leptin action through hypothalamic nitric oxide synthase-1-expressing neurons controls energy balance. Nat. Med. 18, 820–823 (2012).

23. A. Paul et al., Transcriptional Architecture of Synaptic Communication Delineates GABAergic Neuron Identity. Cell. 171, 522–539.e20 (2017).

24. L. Tricoire, T. Vitalis, Neuronal nitric oxide synthase expressing neurons: a journey from birth to neuronal circuits. Front. Neural Circuits. 6, 82 (2012).

25. D. Gerashchenko et al., Identification of a population of sleep-active cerebral cortex neurons. Proc. Natl. Acad. Sci. U. S. A. 105, 10227–10232 (2008).

26. L. Lo, D. J. Anderson, A Cre-dependent, anterograde transsynaptic viral tracer for mapping output pathways of genetically marked neurons. Neuron. 72, 938–950 (2011).

27. Z. a. Knight et al., Molecular profiling of activated neurons by phosphorylated ribosome capture. Cell. 151, 1126–1137 (2012).

28. Z. A. Knight, S. F. Schmidt, K. Birsoy, K. Tan, J. M. Friedman, A critical role for mTORC1 in erythropoiesis and anemia. Elife. 3, e01913 (2014).

29. J. Kim, X. Zhang, S. Muralidhar, S. A. LeBlanc, S. Tonegawa, Basolateral to Central Amygdala Neural Circuits for Appetitive Behaviors. Neuron. 93, 1464–1479.e5 (2017).

30. J. A. Hardaway et al., Central Amygdala Prepronociceptin-Expressing Neurons Mediate Palatable Food Consumption and Reward. Neuron. 102 (2019), p. 1088.

31. I. Kusumoto-yoshida, H. Liu, B. T. Chen, A. Fontanini, A. Bonci, Central role for the insular cortex in mediating conditioned responses to anticipatory cues (2014), doi:10.1073/pnas.1416573112.

32. A. Yiannakas, K. Rosenblum, The Insula and Taste Learning. Front. Mol. Neurosci. 10, 335 (2017).

33. J. M. Bruce et al., Changes in brain activation to food pictures after adjustable gastric banding. Surg. Obes. Relat. Dis. 8, 602–608 (2012).

34. S. Frank, S. Kullmann, R. Veit, Food related processes in the insular cortex. Front. Hum. Neurosci. 7, 499 (2013).

35. P. S. Hogenkamp et al., Higher resting-state activity in reward-related brain circuits in obese versus normal-weight females independent of food intake. Int. J. Obes.. 40, 1687–1692 (2016).

