## Supplementary Information for "A molecularly defined insular cortex --> central amygdala circuit mediates conditioned overconsumption of food"

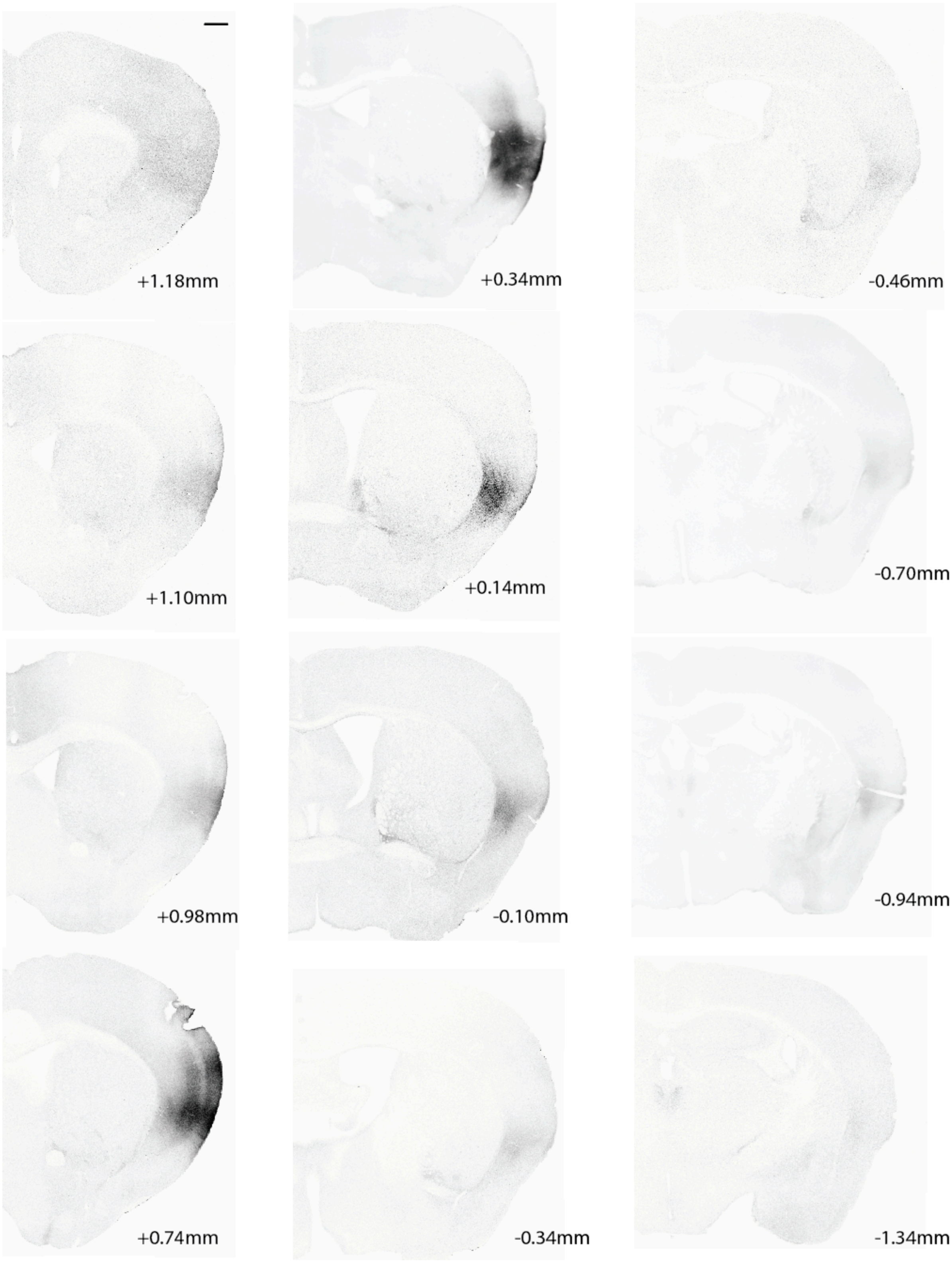
+1.18mm
+0.34mm
-0.46mm
+1.10mm
+0.14mm
-0.70mm
+0.98mm
-0.10mm
-0.94mm
+0.74mm
-0.34mm
-1.34mm

**Fig. S1. Insular cortex injection site.** Representative images of the insular cortex injection site throughout the brain of mice injected with an AAV expressing CaMKIIa-hM4Di-mCherry. Scale bar, 500μm.

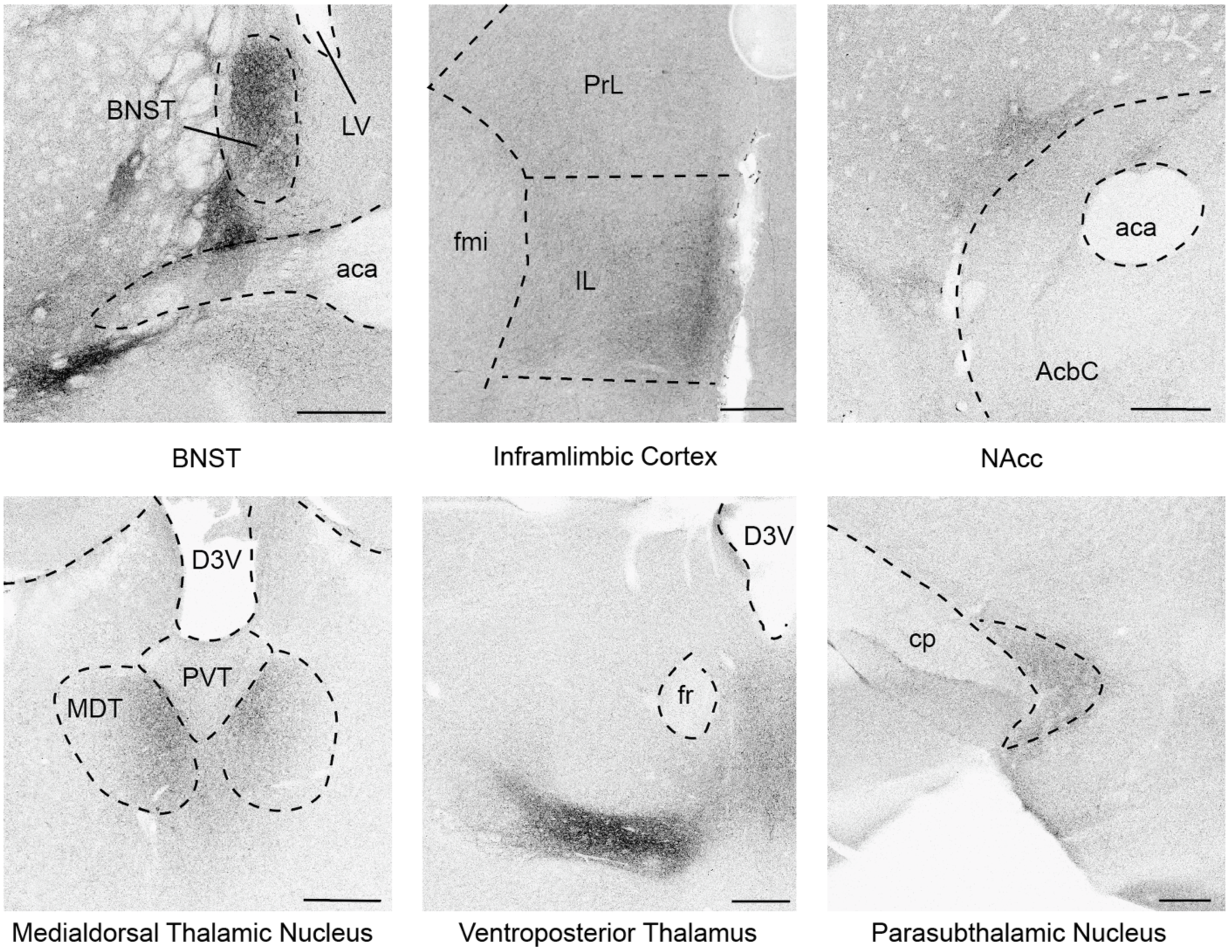


**Fig. S2. Insular cortex projections.**

Mice injected with an AAV expressing CaMKIIa-mCherry into the insular cortex were examined for terminals throughout the brain. Terminals were found in the Bed Nucleus of Stria Terminalis (BNST), Infralimbic Cortex, Nucleus Accumbens (NAcc), Mediodorsal Thalamic Nucleus, Ventroposterior Thalamus and the Parasubthalamic Nucleus. Scale bars, 250μm.

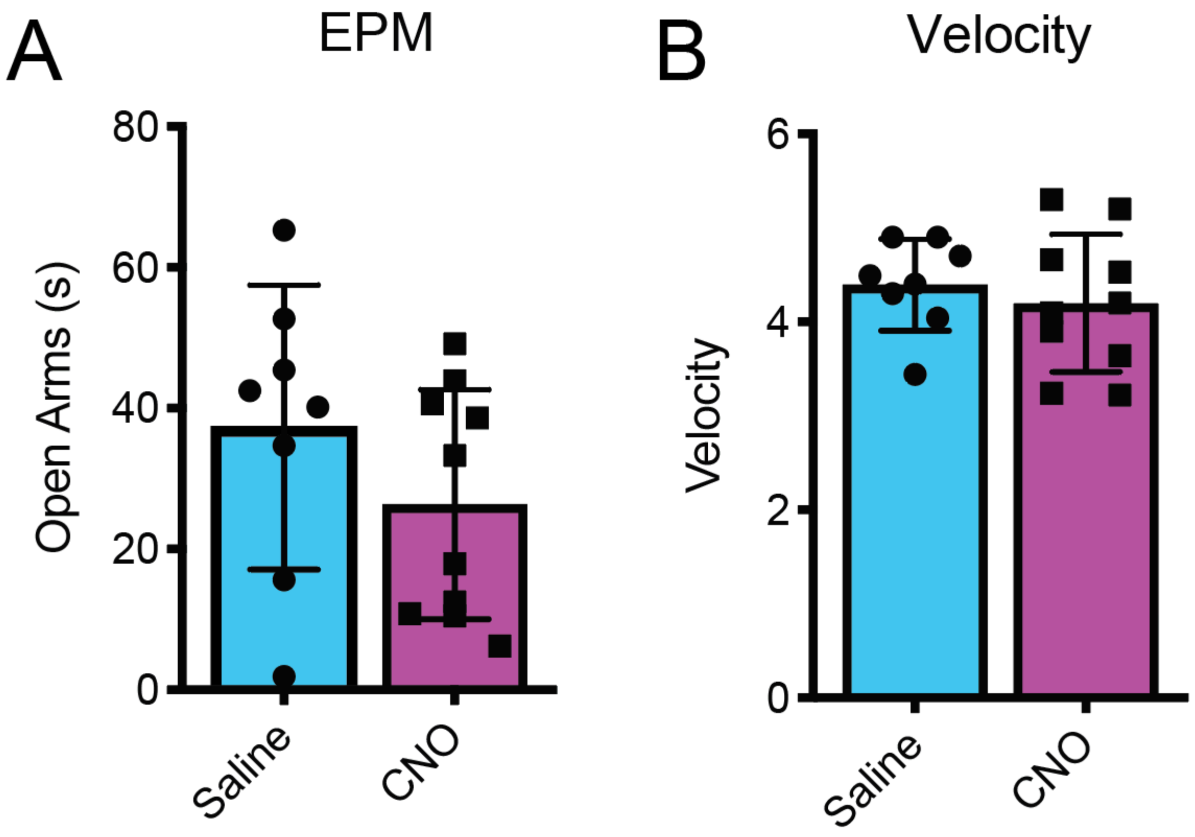


**Fig. S3. Inhibition of insular cortex --> CeA terminals does not effect anxiety or locomotion.** A. Time, measured in seconds, s, spent in the open arms of an elevated plus maze (EPM) (n=8-10). B. Velocity, measured in centimeters transvered per s (cm/s) (n=8-9).

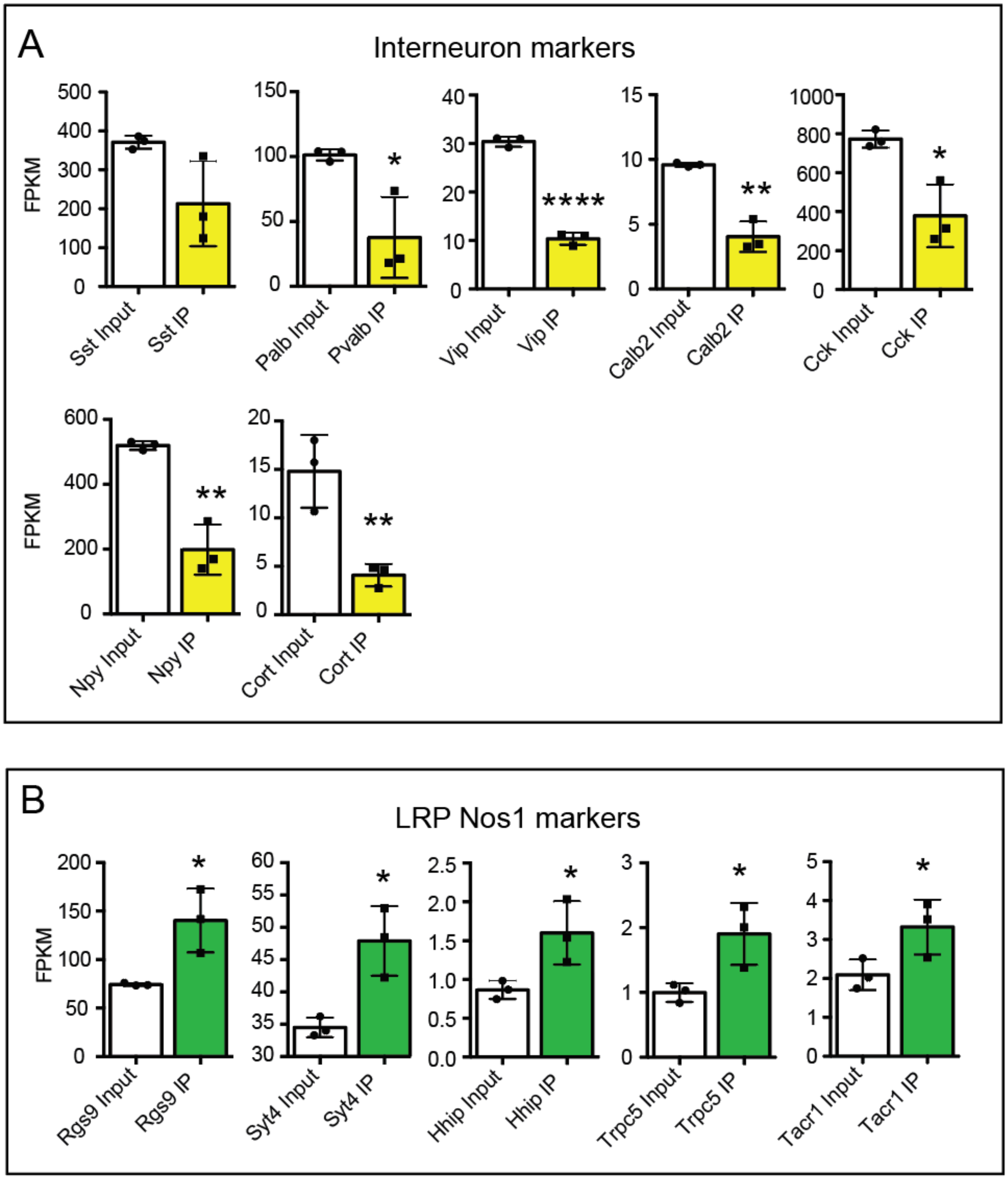


**Fig S4. Analysis of retro-TRAP dataset.** A. Depletion of inhibitory interneuron marker genes in the IP fraction (yellow) compared to the Input (white, n=3, unpaired t-tests. B. Enrichment of long-range projection (LRP) Nos1 marker genes in the IP fraction (green) compared to the Input (white, n=3, unpaired t-tests

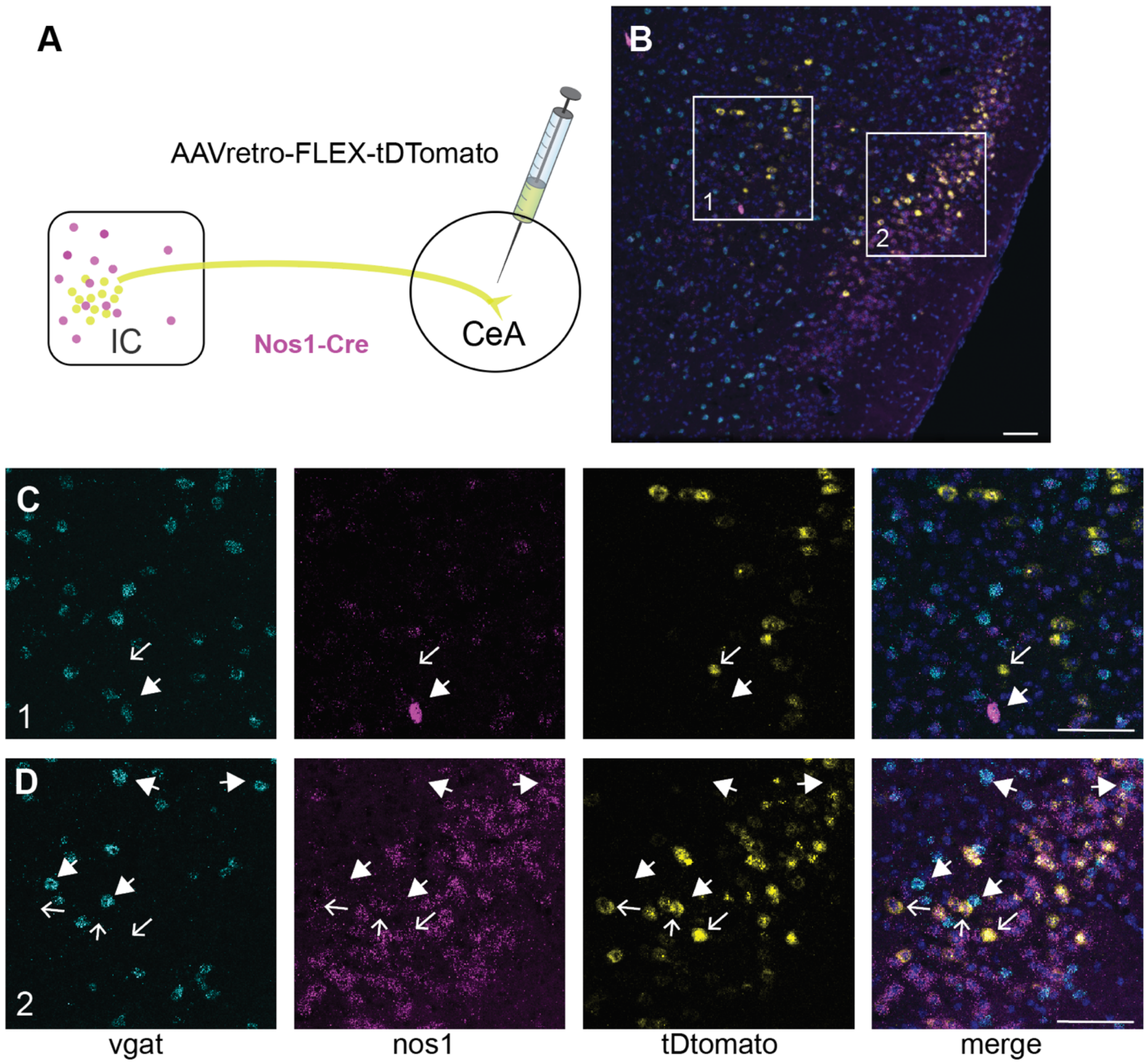


**Fig S5. Nos1-Vgat neurons do not project to CeA.** A. Experimental scheme. Nos1-cre mice were injected with a retrograde AAV expressing tdTomato in a cre-dependent manner. B. In situ hybridization image of the insular cortex. **I**nset squares are shown magnified in C and according to the numbered scheme. C. Vgat (blue), Nos1 (magenta) and tDtomato (yellow) pseudocolored images of Layer 5 neurons in the insular cortex. Closed arrow points to Nos1-Vgat co-expression that is lacking mCherry. mCherry expressing neurons do not express Vgat (open arow points to an example). D. Vgat (blue), Nos1 (magenta) and tDtomato (yellow) pseudocolored images of Layer 2/3 neurons. mCherry expressing neurons co-express Nos1 (open arrows points to examples), whereas Vgat expressing neurons do not co-express mCherry (closed arrows) point to examples. Scale bars, 80um.

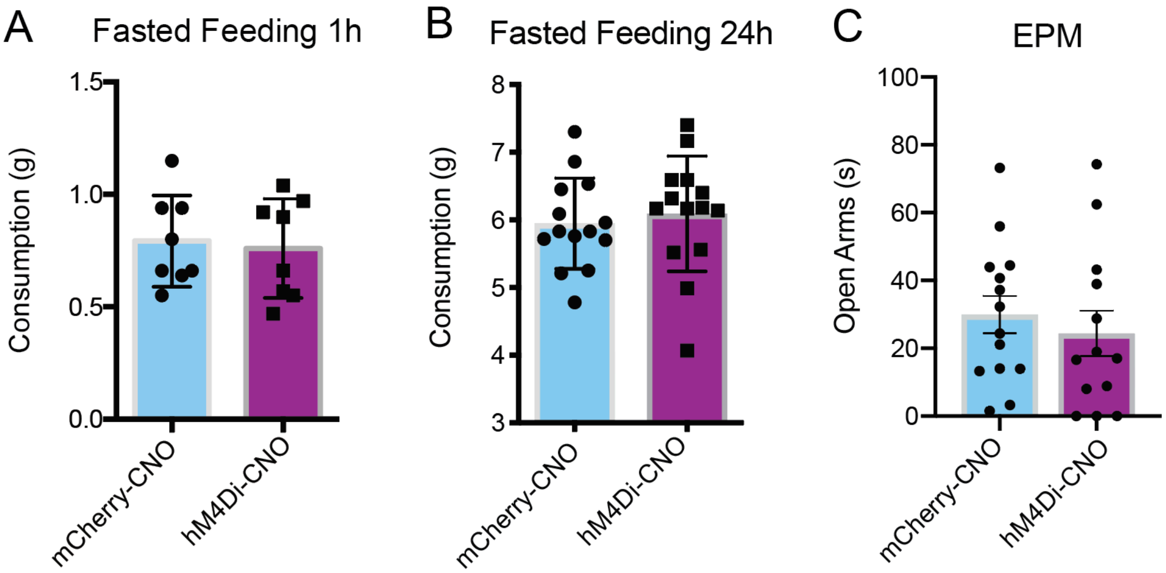


**Fig S6. Inhibition of insular cortex Nos1 neurons does not alter homeostatic feeding or anxiety.** A. Food consumption, measured in grams (g), for 1 hour following an overnight fast (n=8). B. Food consumption, measured in g, over a 24 hour period following an overnight fast (n=14). C. Time, measured in seconds, s, spent in the open arms of an elevated plus maze (EPM) (n=10-11).

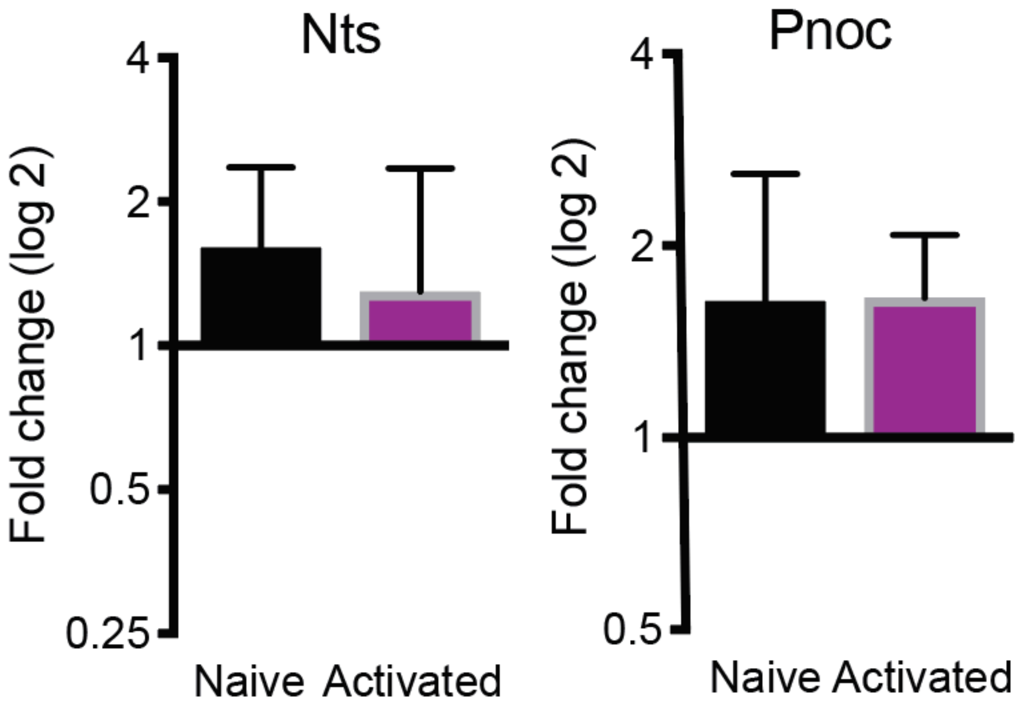


**Fig S7. Nos1 activation does not alter other CeA markers.** Fold-change determined by qPCR of CeA markers, Nts and Pnoc.
